# THE POSSIBILITY OF THE USE OF LEUKOCYTES IN WEAK BLOOD GROUP ANTIGENS DETECTION

**DOI:** 10.1101/2024.02.13.580159

**Authors:** P.G. Kravchun, F.S. Leontyeva, O.D Povelichenko, V.Yu. Dielievska

**Affiliations:** Kharkiv National Medical University; Sytenko Institute of Spine and Joint Pathology National Academy of Medical Sciences of Ukraine

**Keywords:** weak, leukocytes, erythrocytes, antibody, transplantation

## Abstract

**Introduction:** Mixed agglutination reaction is used for revealing A,B and H blood group antigens on the cells.

The aim of the study was to visualize the antibody binding on weak blood group antigens of leukocytes in order to improve the quality of blood typing.

**Material and methods:** The antibodies were contacted with group A and B leukocytes at 4 °C. Mixed agglutination reaction was used to reveal group antigens on leukocytes using test erythrocytes.

**Results:** The heated anti-A,B serum after adsorption on group A leukocytes agglutinated group A erythrocytes.

Anti-A,B heated plasma after the contact with group B leukocytes agglutinated group B erythrocytes.

Anti-A,B serum after adsorption on group B(A+) leukocytes agglutinated group B (A+) erythrocytes.

Anti-A,B serum after adsorption on group A(B+) leukocytes agglutinated group B erythrocytes.

**Conclusions:** MAR allowed to reveal both agglutinogenic and weak adsorbing blood group antigens, that helps to perform accurate blood group typing, especially for transplantation goals.

The study revealed, the detection of adsorbing antigens of leukocytes should be performed at 4 °C.

## Introduction

The leukocytes and erythrocytes have demonstrated the presence of blood group antigens [1,2], however the characteristics of these antigens were reported to be different.

The investigation of the blood group antigens on leukocytes, as nuclear cells, is important and, the possibility for blood group typing using leukocytes is necessary to investigate.

The cells were reported to be coated with IgG for visuaization their antigens.

Thus, the researchers used the latex particles coated with IgG, that attach to the region of the spermatozoa to reveal the attached antisperm antibodies are located. Specific anti-IgG binded to the mixture of fresh spermatozoa and latex beads. There tests for IgA and IgM were also developed [3].

The authors reported of the test with a washed suspension of Rhesus (Rh)-positive human erythrocytes coated with Rh-directed human IgG antibodies, that are are mixed with fresh semen. After the addition of antihuman IgG antibody, IgG containing antisperm antibodies are then binding with the IgG-coated erythrocytes. When spermatozoa are antibody bound, they will form mixed agglutinates with the erythrocytes. The spermatozoa that have adherent particles are calculated [4-7].

The adsorption technique is used in blood group typing for the detection of weak blood group antigens as an indirect method demonstrating the decreased activity of specific antibodies. The fact, that antibodies are specifically adsorbed on the cells stimulated the search of the method of direct visualization of their binding with the cells [8].

The studies reported of different localization of A,B and H blood group substances on erythrocytes and cells [7-9], linked to the membranes of the cells in the glycolipid form or in the composition of glycoproteins in the secretions [10].

The blood group antigens on the cells have been investigated by MAR [11].

The possibility of visualization of adsorbed antibodies on weak blood group antigens of nucleated cells have led us to investigate the method of mixed agglutination for detection of rare blood group types missed by usual agglutination technique.

## Material and methods

The set of monoclonal antibodies was used for forward blood group typing by standard procedure [12].

The samples of citrated plasma, EDTA plasma and serum were used as a source of antibodies for MAR.

The leukocytes were taken from EDTA blood and washed with normal saline.

The leukocytes were centrifuged for one minute at 1000 rpm. The supernatant fluid was removed and the deposited cells were resuspended in the saline.

The erythrocytes from EDTA blood were washed and used as test group A and B erythrocytes. 0.5% suspension of erythrocytes was mixed with diluent: group AB plasma or serum (1:100 in saline).

The samples of anti-A,B plasma from the persons with O blood group type were heat-inactivated for 30 minutes at 56 °C before the use to obtain IgG group specific antibodies.

### Mixed agglutination reaction

The study was performed with leukocytes and erythrocytes by the method described by Coombs [13-15].

50 microliters of group B leukocytes was incubated with 50 microliters of anti-A,B serum, citrated or heated anti-A,B plasma and 50 microliters of normal saline at 4 °C for 13 hours. The supernatant fluid was analyzed with group B erythrocytes on the presence of agglutination after adsorption.

The cells after adsorption were centrifuged for two minutes at 1000 rpm, washed three times in saline and group A and B erythrocytes were added in AB serum or plasma (1:100).

The mixtures were incubated for one hour at 4°C and at room temperature, centrifugated for two minutes at 1000 rpm and the samples were investigated under microscope. Statistical analysis was performed by Statistica 10.0 software program.

The study was approved by Ethical committee of Kharkiv National Medical University (protocol N 2). Informed consent was obtained from the participants with O, A and B blood group persons. The study used convenience sampling method.

## Results

### The detection of group A antigen on leukocytes

Anti-A,B serum after adsorption on group A(B+) leukocytes agglutinated A(B+) erythrocytes.

Anti-A,B serum after adsorption on group A leukocytes agglutinated group A erythrocytes and did not agglutinate B erythrocytes.

The heated anti-A,B serum after adsorption on group A leukocytes agglutinated group A erythrocytes in two samples (Figures 1). Another sample of anti-A,B serum after adsorption on group A leukocytes agglutinated group A erythrocytes.

**Figure 1.**
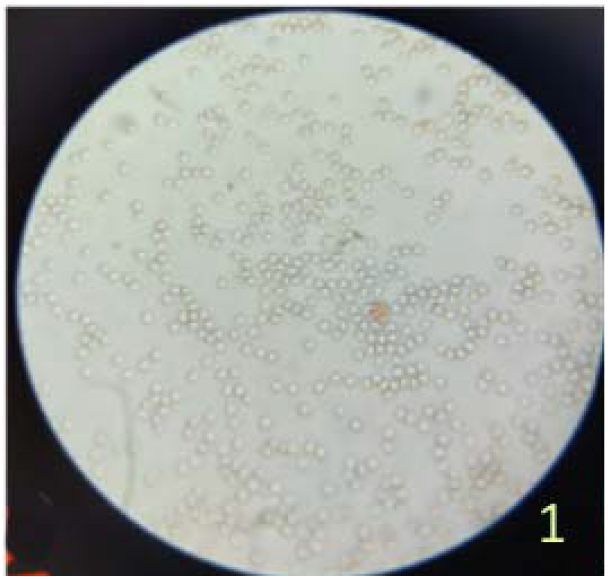
The heated anti-A,B serum after adsorption on group A leukocytes agglutinated group A erythrocytes.

Anti-A,B serum after adsorption on group A leukocytes agglutinated group A erythrocytes (Figure 2).

**Figure 2.**
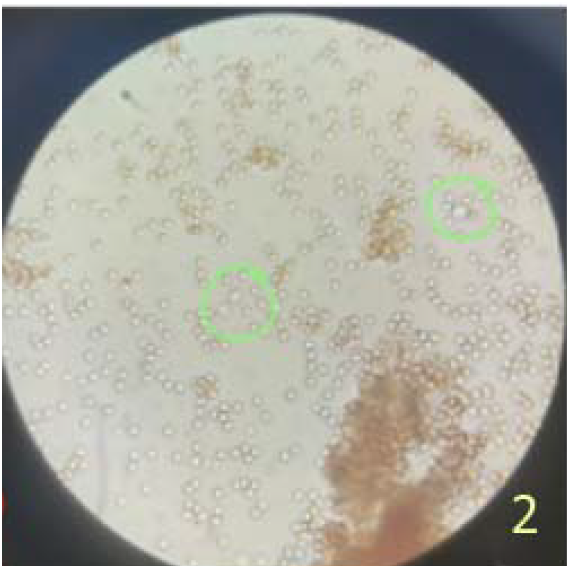
Anti-A,B serum after adsorption on group A leukocytes agglutinated group A erythrocytes.

Anti-A,B heated plasma after adsorption on group A leukocytes agglutinated group A erythrocytes.

Anti-A,B citrated plasma after adsorption on group A(B+) leukocytes agglutinated group A(B+) erythrocytes.

### Detection of group B antigen on leukocytes

Anti-A,B heated plasma after adsorption on group B (A+) leukocytes agglutinated group B erythrocytes in two samples.

Anti-A,B heated plasma after the contact with group B leukocytes agglutinated group B erythrocytes (Figures 3, 4).

**Figure 3.**
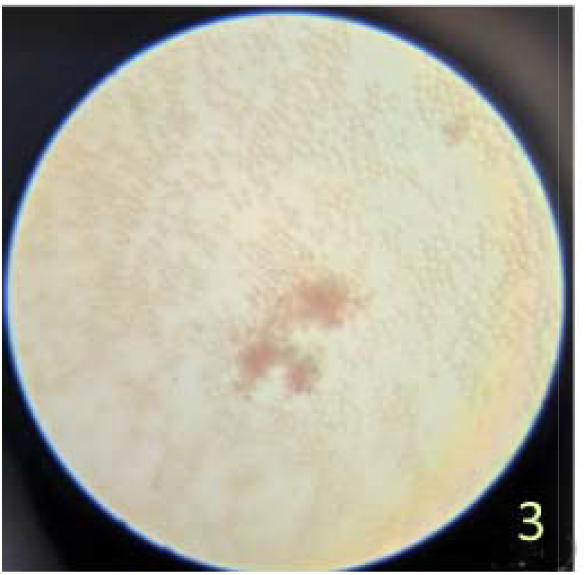
Anti-A,B heated plasma after the contact with group B leukocytes agglutinated group B erythrocytes.

**Figure 4.**
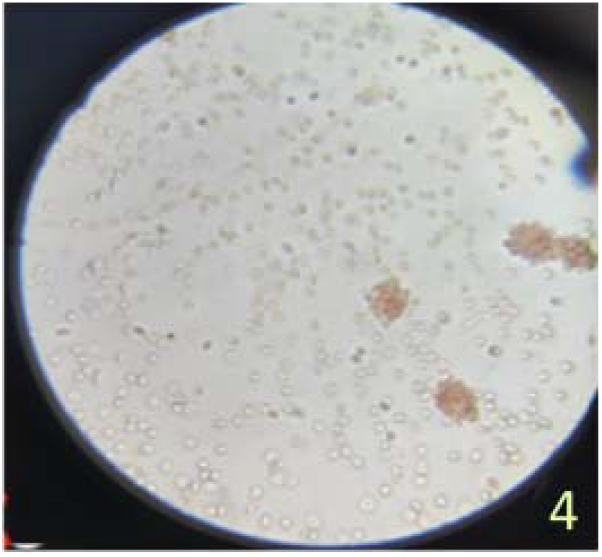
Anti-A,B heated plasma after the contact with group B leukocytes agglutinated group B erythrocytes.

### Detection of weak group A antigen on leukocytes

Anti-A,B heated serum after adsorption on group B(A+) leukocytes attached group A erythrocytes (Figure 5). In two samples anti-A,B serum being adsorbed on group A leukocytes aggutinated group B(A+) erythrocytes (Figure 6). Anti-A,B heated plasma after adsorption on group B(A+) leukocytes agglutinated group A erythrocytes. Anti-A,B heated plasma after adsorption on group A leukocytes agglutinated group A erythrocytes.

**Figure 5.**
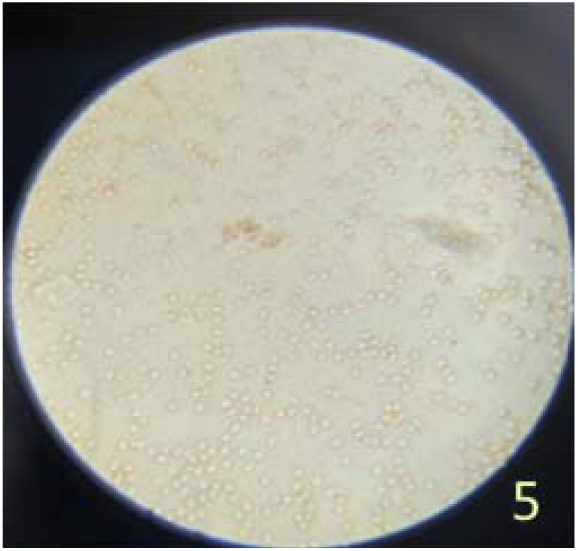
Anti-A,B heated serum after adsorption on B(A+) leukocytes attached B (A+) erythrocytes.

**Figure 6.**
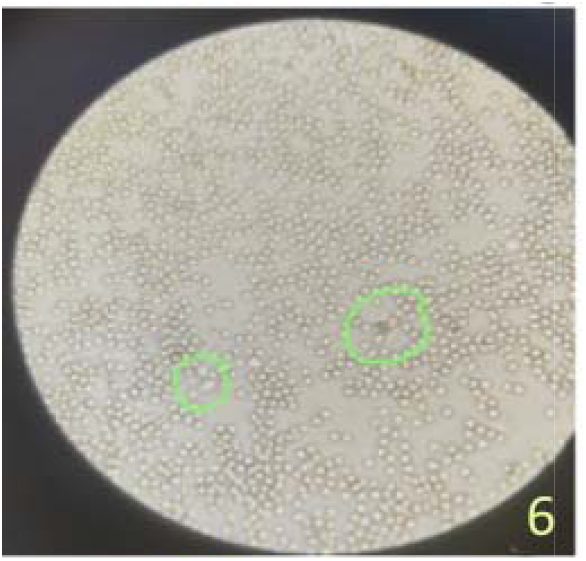
Anti-A,B serum being adsorbed on group A leukocytes aggutinated group B(A+) erythrocytes.

### Detection of weak adsorbing group B antigen on leukocytes

Anti-A,B serum after adsorption on group A(B+) leukocytes attached group B erythrocytes (Figure 7).

**Figure 7.**
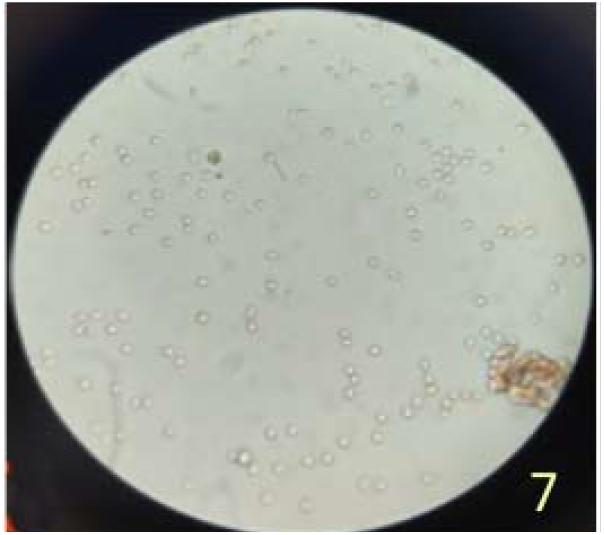
Anti-A,B serum after adsorption on group A(B+) leukocytes attached B erythrocytes.

Anti-A,B serum after adsorption on group A(B+) leukocytes agglutinated group B erythrocytes (Figure 8).

**Figure 8.**
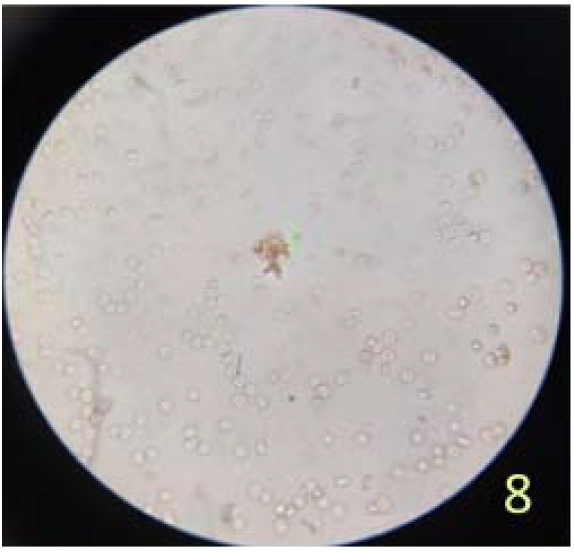
Anti-A,B serum after adsorption on group A(B+) leukocytes agglutinated group B erythrocytes.

### Absence of agglutination in mixed aglutination reaction

Anti-A,B EDTA plasma after adsorption on group O leukocytes did not agglutinate group B erythrocytes (at 4°C and 37°C).

Anti-A,B plasma after adsorption on group B leukocytes did not agglutinate group A erythrocytes (Figure 9).

**Figure 9.**
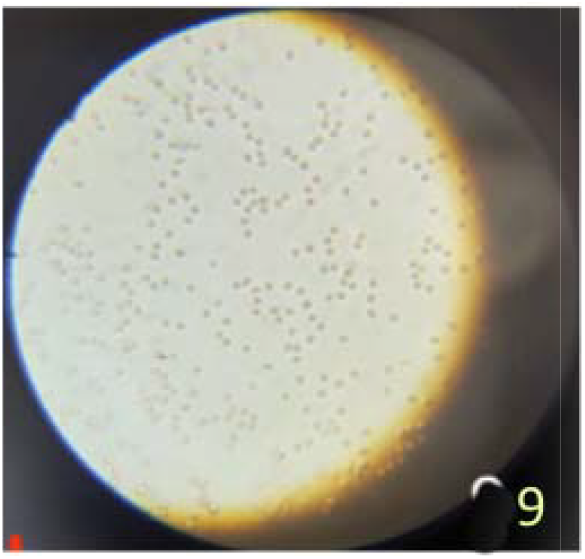
Anti-A,B plasma after adsorption on B leukocytes did not agglutinate A erythrocytes.

Anti-A,B heated serum after adsorption on group A leukocytes did not agglutinate group B erythrocytes (Figure 10).

**Figure 10.**
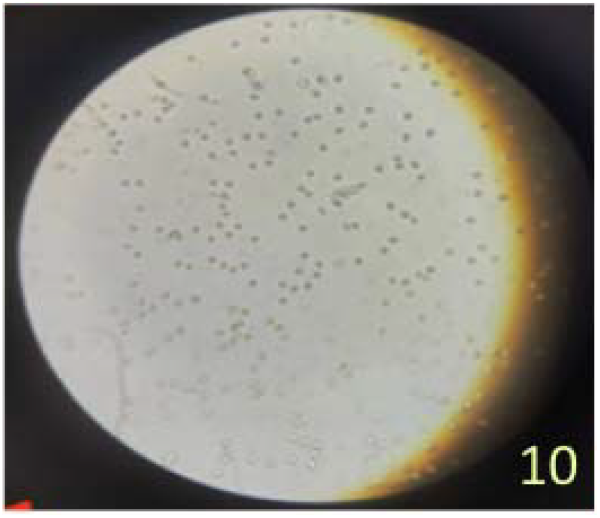
Anti-A,B heated serum after adsorption on group A leukocytes did not agglutinate group B erythrocytes.

Anti-A,B serum after adsorption on group A(B+) leukocytes did not agglutinate group A (B+) erythrocytes. Anti-A,B serum after adsorption on group A leukocytes did not agglutinate group A erythrocytes. Anti-A,B heated serum after adsorption on group A leukocytes did not agglutinate group A erythrocytes, however another sample of the serum (nonheated) after adsorption on group A leukocytes agglutinated group A erythrocytes. The sera were not investigated on the titer and characteristics of antibodies.

### The use of monoclonal antibodies in MAR

Monoclonal antibody 2-10 after adsorption on group A leukocytes agglutinated group A erythrocytes (Figure 11).

**Figure 11.**
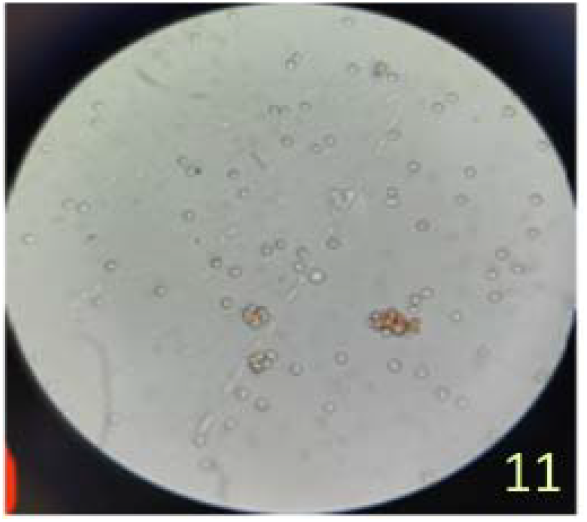
Monoclonal antibody 2-10 after adsorption on group A leukocytes agglutinated group A erythrocytes.

Monoclonal anti-A antibody after adsorption on group A leukocytes agglutinated group A erythrocytes and did not agglutinate group B erythrocytes.

Thus, the persons typed as blood group A by method of erythrocytes agglutination demonstrated the presence of group A antigen on leukocytes by mixed agglutination reaction.

Importantly, MAR revealed the presence of weak group A antigen, detected by adsorption test and being not revealed by usual agglutination reaction.

In persons typed as blood group B by the method of erythrocytes agglutination group B antigen was determined on leukocytes by MAR.

In persons typed as blood group O by the method of erythrocytes agglutination the leukocytes did not show the presence of group A or B antigens by MAR.

Thus, MAR allowed to reveal not only agglutinogenic, but also adsorbing weak blood group antigens, that helps to perform accurate blood group typing, especially for transplantation goals.

The study revealed better detection of adsorbing antigens on leukocytes at low temperature: adsorption was performed at 4 °C, the binding of test erythrocytes to adsorbed antibodies on leukocytes was performed at 4°C and 23 °C.

The contact of anti-A,B heated serum with leukocytes in diluent caused less expressed agglutination of test erythrocytes as compared to the contact with leukocytes in saline. The contact of anti-A,B heated serum with normal saline (1:1) and leukocytes led to the less but more sensitive agglutination with erythrocytes as compared to the serum without normal saline. Heated anti-A,B serum after the contact with group A leukocytes agglutinated group A erythrocytes more significantly as compared to the usual anti-A,B serum. Thus, the test erythrocytes in diluent were contacted with sensitized leukocytes.

To estimate the possibility of the use of any polyclonal serum in mixed agglutination reaction for group leukocyte antigens detection the anslysis of different sera was performed in MAR.

Thus, anti-A,B serum of the person without anti-A antibody adsorbing ability of erythrocytes after the contact with group A leukocytes agglutinated two samples of group A erythrocytes and B (A+) erythrocytes. However, anti-A,B serum of another sample of the person with anti-A antibody adsorbing ability of erythrocytes after the contact with group A leukocytes agglutinated only one sample of group A erythrocytes and did not agglutinate another sample of group A erythrocytes and B (A+) erythrocytes. The anti-A antibodies from the serum from the person from O blood group person without anti-A antibody adsorbing ability showed better adsorption on group A leukocytes, whereas anti-A antibodies from O blood group person with anti-A adsorbing ability showed better adsorbing ability on group A erythrocytes.

Thus, before the use of polyclonal serum for MAR the erythrocytes of the person should be investigated on adsorbing ability of anti-A and anti-B antibodies. Absence of anti-A and anti-B adsorbing ability of erythrocytes in O blood group persons testifies to the possibility of the use of their serum.

The distinctions between the described MAR and the method used in the study.

1. The leukocytes were used instead of epithelial cells.
2. The adsorption reaction was performed for 13 hours (not for 1 hour)
3. The adsorption reaction was performed at 4 °C (not at 23 °C).
4. The serum, citrated plasma and EDTA plasma were investigated for comparison and the best reproducible results were obtained with serum.
5. Different adsorbing abilities of antibodies were revealed in blood group O sera, depending on the characteristics of containing antibodies, their titer and optimal temperature of the reactivity.

**Table 1.** The results of mixed agglutination reaction in revealing group A and B agglutinogenic and adsorbing antigens.

|  | adsorption<br>on<br>leukocytes | positive<br>reaction with<br>erythrocytes | negative<br>reaction with<br>erythrocytes |
| --- | --- | --- | --- |
| anti-A,B<br>serum | A(B+) | A(B+) | B |
| anti-A,B<br>serum | A | A |  |
| anti-A,B<br>heated serum | A | A |  |
| citrated<br>anti-A,B<br>plasma | A(B+) | A(B+) |  |
| heated anti-A,B plasma | A | A |  |
| anti-A,B<br>serum | A(B-) | B(A+) |  |
| heated anti-<br>A,B serum | B(A+) | A |  |
| anti-A<br>monoclonal<br>antibody | A | A | B (A+) |
| anti-A<br>monoclonal<br>antibody 2-10 | A | A |  |
| anti-A,B<br>serum | A(B+) |  | A(B+) |
| anti-A,B | A |  | A |
| serum |  |  |  |
| heated anti-A,B serum | B(A+) | B |  |
| heated anti-A,B serum | B | B |  |
| anti-A,B serum | A(B+) | B |  |
| citrated anti-A,B plasma | A(B+) | B |  |
| EDTA anti-A,B plasma | O |  | B at 4 °C<br>at 37 °C |

## Discussion

MAR with immune anti-A sera was reported to reveal group A antigen on the cells of group A, A2 and group A1 persons [16]. The strength of the reaction with the cells was not associated with the content of group A antigen in the saliva as measured by the agglutination inhibition of the serum [10,17].

The ultrastructural localization of blood group substance A and its relationship with the cell coat was studied on exfoliated buccal epithelial cells. Group A antigen was detected by a double layer immunoperoxidase staining method and a double layer immunoferritin staining method.

MAR was reported to detect A,B and H blood group antigens on buccal cells [11, 18].

Group B and H isoantigens of human gastric and small intestinal mucosae detected by MAR, were absent in eight of nine gastric epithelial malignancies.

Different methods for the detection of weak group antigens have been reported [19]. The studies showed an importance of revealing of weak group antigens for safe blood transfusions [16-18].

Group A,B and H antigens cause an IgM immune response with greatest reactivity at 4 °C and are present in nature. This exposure is the basis for the production of “naturally occurring” antibodies (anti-B antibodies in A blood group).

Compared with manual methods, automated methods for blood group typing have been expected to be sufficient [20, 21]. However, automated methods were reported to demonstrate mixed fields, specimen turbidity, mismatch results or weak reactions and manual methods are used to confirm ABO grouping [21, 22].

The use of antiglobulin-based tests have been accepted for the detection of the antibodies bound to the surface of the cells, including the mixed anti-globulin reaction test. The tests, that reveal the percentage of antibody-coated motile spermatozoa, were recommended by WHO as screening tests for all the semen samples examined in the couple-infertility work-up (WHO, 2010), considering 50% antibody-coated motile spermatozoa as the clinically-relevant threshold.

The present study intended to visualize antibody binding with weak blood group antigens and to reveal the optimal conditions of their interaction in MAR using leukocytes and erythrocytes for improvement of blood group typing.

The majority of the researchers, including R. Coombs reported of the use of the heated serum for the study of adsorbing antibodies [9]. The heated plasma is known to contain IgG antibodies, and adsorption of IgG antibodies on leukocytes corresponds to the notion of high avidity of IgG antibodies to the cells on the contrary to IgM antibodies. Many anti-A sera were reported to be very poor at producing MAR despite adequate hemagglutinating properties. The different abilities of producing mixed agglutination of the sera from individuals were found to be related to the presence of 7S or 19S fractions and presence of hemolysins [24].

Incomplete blood group antibodies were reported to be 7S globulins, agglutinating antibodies were 19S globulins. Incomplete antibodies were diminished by the treatment of the serum with 2-mercaptoetanol. 19S globulins with agglutinating activity and 7S globulins with ability to sensitize erythrocytes to an antiglobulin serum evidenced of two kinds of antibodies. 7S antibodies are known to be presented by IgG antibodies, 19S antibodies are reported to compose IgM antibodies. IgM and IgG antibodies are known to compete for the same substrate antigen. The data indicate that passively administered antibody has depressive effect on immune response. The evidence suggests that antibody inhibits the response by combining with antigenic determinants [25,26].

Pearlman reported that both 19S and 7S antibobodies are effective modificators of antibody formation both are capable of enhancement or depression of immunologucal response when transfered in small quantities [27,28].

The conducted study revealed the possibility of the use of MAR in detection of weak blood antigens at 4°C with the use of the polyclonal serum or heated polyclonal serum instead of monoclonal antibody.

## Conclusions

MAR allowed to reveal both agglutinogenic and weak adsorbing blood group antigens, that helps to perform accurate blood group typing, especially for transplantation goals.

The study revealed, the detection of adsorbing antigens of leukocytes should be performed at 4 °C with prolonged adsorption period.

